# Utility of Kirschner Wires as Prosthetic Implants to Investigate Establishment and Treatment of Biofilm-Associated Infections in a *Galleria mellonella* Model

**DOI:** 10.64898/2026.02.26.708268

**Authors:** Tia Arnaud, Natasha Theriault, Deborah Ann Court, Steven Serge Theriault, Tasia Joy Lightly, Bradley William Michael Cook

## Abstract

**Objectives:** Bacteriophage (phage) therapy is being explored as a strategy for clinical control of periprosthetic joint infections, addressing the challenges of antibiotic-resistant bacteria and limitations around the complete eradication of methicillin-resistant *Staphylococcus aureus* biofilms by conventional antibiotic treatment. Additionally, phage-antibiotic combinations may enhance treatment success through synergistic interactions. This study aimed to compare the efficacy of phage *Silviavirus remus* (Remus) and vancomycin, independently and together, against *S. aureus* biofilms on Kirschner-wire (K-wire) implants using *Galleria mellonella* as an *in vivo* model, and to assess the biocompatibility of K-wire implantation.

**Results:** Survival of *G. mellonella* larvae implanted with *S. aureus* biofilms and treated with Remus (∼3 × 10^4^ PFU/worm) and/or vancomycin (5 µg/worm) after 72 hours was assessed using Kaplan-Meier curves. Statistical analyses among treatment groups were performed using log-rank tests at p-value < 0.10. Implantation of uncolonized K-wires with treatment administration did not affect survival (100%). Compared with untreated biofilm-infected larvae (77%), vancomycin alone (97%; p = 0.023) or in combination with Remus (93%; p = 0.087) improved survival, whereas Remus alone did not increase survival (67%, p = 0.33). Scanning electron microscopy confirmed the presence of biofilm-associated attachment of *S. aureus* on K-wires, although adherence was non-uniform.

## Introduction

Periprosthetic joint infections (PJIs) remain difficult to eradicate with conventional antibiotic therapy, largely due to the ongoing surge of antibiotic resistance and biofilm formation on the surfaces of orthopedic prostheses, predominantly by methicillin-resistant *Staphylococcus aureus* (MRSA). As a result, bacteriophage (phage) therapy is being explored as an alternative treatment strategy to overcome these challenges.

*Silviavirus remus* (Remus) is a strictly virulent staphylococcal phage with bacteriolytic and anti-biofilm activity against *S. aureus* [1]. Previous *in vitro* application of Remus showed its ability to prevent and remove biofilms when used as a coating on ceramic implants, demonstrating its potential use against PJI-associated infections [2]. However, to our knowledge, this study will be the first exploration of its efficacy against implant-associated biofilms *in vivo*.

*G. mellonella* larvae have previously been used as a model to assess the biocompatibility and utility of stainless-steel K-wires for inoculation and treatment of preformed biofilms by implantation [3]. While a biofilm infection model was successfully established *in vivo*, treatment with gentamicin did not rescue larvae [3]. Similar work demonstrated the utility of K-wires to investigate the efficacy of phage Sb-1 against *S. aureus* biofilms on indwelling devices [4]. Although Sb-1 alone reduced K-wire colonization to comparable levels of vancomycin, its efficacy was markedly improved when combined with daptomycin [4].

The current study aimed to further explore the use of K-wires in assessing phage-antibiotic therapy against PJI-associated biofilms using a *G. mellonella* model. The data presented support previous reports of biocompatibility and provide survival outcomes following phage and antibiotic individual and combined therapies. This research note supports the utility of this underutilised, cost-effective method, which can be extended to other implant-associated infections. Conducted as a short pilot study to test feasibility, this work was not included in a larger manuscript due to its preliminary nature.

## Materials and Methods

### Bacteria strains and bacteriophage

Methicillin-resistant *Staphylococcus aureus* (MRSA) PS47 and phage *Silviavirus remus* (Remus) were purchased from the Felix d’Hérelle Reference Center for Bacterial Viruses (University of Laval, Quebec City, Canada). *S. aureus* PS47 was used for routine propagation of Remus. An isolate of *S. aureus* capable of biofilm formation, previously described [5], was used for biofilm formation on stainless-steel implants. Bacterial strains were routinely cultured in Tryptic Soy Broth (TSB) or on Tryptic Soy agar (TSA) at 37°C for 18 hours, unless otherwise stated.

### Bacteriophage propagation, concentration, and purification

Phage Remus was propagated, concentrated and purified as described previously [5]. Once purified, Remus was titrated for plaque-forming-units (PFU)/mL on TSA soft agar overlay, according to methods outlined previously [6], with PFUs determined after overnight incubation at 37°C.

### *In vivo* bacterial infection and treatment of *Galleria mellonella* larvae

*G. mellonella* larvae were obtained from Serum Therapeutics Inc. (Edmonton, Canada). Upon delivery, larvae were maintained in the dark on-feed at 24°C. At least one day prior to use, larvae were removed from feed and starved at 24°C. Groups of ten larvae (fifth – sixth instar) were selected based on similar weight (≥ 230 mg), and lacking signs of pupation or melanization.

### Establishment of biofilms on K-wires

Stainless-steel K-wires (0.8 mm x 4-6 mm; BlueSao Canada, Laval, Canada) were used for implantation of biofilms in *G. mellonella*. Briefly, an overnight culture of *S. aureus* was adjusted to 1 × 10^6^ colony-forming-units (CFU)/mL in TSB with 0.5% (w/v) glucose. Five sterile K-wires were placed together in 500 µL of this suspension and incubated at 37°C with shaking at 150 rpm for 24 hours to encourage biofilm formation. Control K-wires (biofilm-free) were prepared in TSB with 0.5% (w/v) glucose without bacteria. K-wires were gently washed twice with equal volumes of sterile phosphate-buffered saline (PBS) to remove planktonic bacteria. To enumerate biofilm-embedded bacteria: three K-wires were selected as representative samples and each placed in 1.7 mL tubes containing 100 µL of sterile PBS. Bacteria were dislodged by sonication (Symphony Ultrasonic Cleaner, VWR, Mississauga, Canada) for 3 minutes at 35 kHz and agitation with a vortex for 1 minute. The sonication fluid was then serially-diluted, 10-fold in PBS and titrated on TSA for 24 hours at 37°C. Bacterial colonies were counted and reported as CFU/K-wire.

### Establishment and treatment of infection in *G. mellonella*

Implant-associated infections were established by inserting K-wires with preformed biofilms (target dose: 3 × 10^3^ CFU) between the second and third proleg segment of larvae, previously anesthetized for 10 minutes on-ice [7]. Infection control groups were implanted with biofilm-free K-wires. Larvae in the planktonic control group were injected in the left hind proleg with 10 µL of 3 × 10^5^ CFU/mL using a CarePoint^™^Vet 31-gauge, 3/10-cc insulin syringe (Allison Medical Inc., Littleton, USA), matching the intended implant bacterial dose. Approximately 1-hour post-inoculation, larvae were treated with 10 µL of Remus (3 × 10^6^ PFU/mL per worm) and/or vancomycin (5 µg per worm); or saline-magnesium (SM) buffer and/or sterile distilled water for treatment control groups. The treatments were administered sequentially, first in the right proleg and then in the second-last right proleg. Larvae were incubated at 37°C and monitored daily for survival for 72 hours post-treatment by unresponsiveness to stimuli. Three biological replicates were performed with 10 larvae per group. The injected doses of planktonic bacteria and Remus were verified by titrating for CFUs and PFUs, respectively, as described above.

### Fixation and SEM imaging of biofilm-associated attachment on K-wires

Biofilm-associated attachment was established on three K-wires, as described above. Negative control wires were also prepared by incubating in uninoculated media. Fixation was performed following methods previously described [3,8]. Briefly, K-wires were submerged in 2% (v/v) glutaraldehyde and incubated overnight, statically at 4°C. Glutaraldehyde was removed, and wires were washed 3 times in PBS and sequentially dehydrated in the following ethanol concentrations (v/v): 10%, 30%, 50%, 70%, 90% and 95%. Residual ethanol was evaporated overnight, and samples were sputter-coated with gold and palladium (Denton Desk II Sputter Coater) then imaged in a FEI Talos F200X S/TEM at 10 kV.

### Statistical analysis

Statistical analysis was performed using GraphPad Prism version 9.4.1. (GraphPad™ Software Inc., La Jolla, USA). Kaplan-Meier survival curves were used to evaluate daily survival rates of larvae for 72 hours after receiving Remus and/or vancomycin treatment, or no treatment. Statistical significance was calculated between treatment groups by log-rank tests using p-value < 0.10.

## Results

### Survival analysis of *G. mellonella* larvae following implant-associated infection

Following *S. aureus* implant-associated infection and treatment with phage Remus and/or vancomycin, Kaplan-Meier survival analysis was performed to determine daily larval survival over 72 hours (data files 1 and 2). Beginning with control groups, the data shows that after 72 hours, implantation with uncolonized K-wires alone (97%; red diamonds) or with phage-vancomycin treatments (100%; black box) did not negatively impact survival (data file 1). It should also be noted that one mortality was reported in the implanted and untreated group; although, observation suggested this was an implantation error that appeared to cause irrecoverable internal damage. Comparing sham-treated larvae, *S. aureus* biofilm-implants (teal circles) and planktonic (non-biofilm) inoculations (yellow triangles), revealed a greater decrease in larval survival in the biofilm-inoculated group (77% versus 87%, respectively) after 72 hours, albeit not significantly (p = 0.35) (data file 1). When compared to untreated larvae (teal circles; 77%), the addition of vancomycin or the Remus-vancomycin combination treatment appeared to significantly increase survival to 97% (purple cross; p = 0.028) and 93% (green circle; p = 0.087), respectively, after 72 hours, whereas treatment with Remus alone (pink box; 67%; p = 0.33) did not improve survival of larvae (data file 1). Enumeration data verifying challenge doses of *S. aureus* on K-wires are provided in data file 3.

### Scanning electron microscopy

The presence of *S. aureus* biofilm-associated attachment on the surface of the K-wires after 24 hours of incubation was evaluated by SEM (data file 4). Imaging verified attachment by *S. aureus* on K-wires, and shows that biofilm formation was not uniformly distributed across the implants. As expected, no adherence was observed on the uninoculated, negative control implants (data not shown).

## Discussion

The treatment of periprosthetic joint infections remains challenged by antibiotic resistance and biofilm formation on orthopedic implants, prompting investigation of phage therapy as an alternative or adjunct to conventional antibiotics. Moreover, the assessment of phage-antibiotic therapies using *in vivo* models can aid in preliminary evaluations of efficacy and treatment outcomes. The data presented herein demonstrates the power of this invertebrate model as a cost-effective method to mimic implant-associated infections and evaluate phage-antibiotic treatment *in vivo*.

Through Kaplan-Meier survival analysis, implantation of biofilm-free K-wires alone or with Remus-vancomycin treatment did not negatively impact survival, indicating that neither the implants nor treatments were toxic to larvae, as reported elsewhere [3]. In the current study, the one mortality in the implanted-untreated group was attributed to an implantation error rather than metal toxicity.

Biofilm-infected larvae exhibited lower survival than larvae infected with planktonic cells, despite the actual bacterial dose delivered on the implants being lower than the intended dose equivalent to planktonic infection (data file 3). This may reflect the increased resistance of biofilm-embedded bacteria, in comparison to their planktonic counterparts, to host immune responses, as commonly observed on implanted medical devices in humans [9,10]. Nonetheless, a significant increase in survival was observed with the administration of vancomycin alone or in combination with Remus, in comparison to untreated biofilm-infected larvae. Interestingly, the survival of infected larvae treated with Remus alone was not significantly different than untreated larvae. This may be attributed to the biofilm-degrading activity of Remus [1], resulting in the dispersal of bacteria and infections at secondary sites and/or systemically, and underscoring the importance of phage-antibiotic combination therapies to enhance treatment outcomes [11].

SEM analysis was performed to verify adherence of *S. aureus* to the K-wires. Although adherence was confirmed, uniform distribution of *S. aureus* was not present across the implants. Additionally, inconsistent biofilm formation was reflected by the varying CFU counts recovered from the K-wires across replicates. A previous study demonstrated strain-specific differences in *S. aureus* biofilm formation under static and dynamic conditions [12]. It is therefore possible that the conditions adopted for this study limited uniform biofilm development, and require further optimization to establish a more tenacious and consistent biofilm in future studies.

## Conclusion

This study demonstrated that *G. mellonella* larvae, implanted with stainless-steel K-wires, can be used as a practical and cost-effective invertebrate model to evaluate phage-antibiotic therapies against implant-associated infections. Using this model, vancomycin significantly improved the survival of biofilm-infected larvae, both alone and in combination with Remus, whereas Remus alone did not improve survival. Our study also identified challenges in establishing consistent biofilm formation on K-wires, which may be addressed in future studies through initial optimization experiments to improve biofilm formation for subsequent antibiofilm studies. Overall, treatment outcomes from this study highlighted the ability to evaluate phage-antibiotic therapies using this underutilized, invertebrate model.

## Limitations

- Biofilm formation was inconsistent on K-wires after 24-hour incubation according to the enumeration data. Determining the conditions required for a given bacterial strain to colonize, adhere, and form a mature biofilm must be established prior to adopting anti-biofilm testing methods in *G. mellonella*.
- Post-implantation, larvae are as mobile as un-implanted worms and may push out the K-wires, potentially compromising successful establishment of biofilm-related infections. The entirety of the K-wire must be implanted inside the length of the larvae to ensure a consistent challenge dose of biofilms. Initial worm size, weight, and frequent visual scoring is important to ensure that the K-wires remain implanted. Careful implantation is required to avoid unintentional injury and mortality.
- Biofilm-degrading and lytic capabilities can vary between phages and their bacterial targets, which may influence efficacy and treatment outcomes. Therefore, it is recommended to evaluate the antibiofilm and lytic activity, as well as the dosage, of the phage of interest prior to adopting this method.
- In combination therapies, it is challenging to determine the timing of antibiotic administration following the application of biofilm-dispersing phages to mitigate infection spread and increase treatment success. A simple invertebrate model, such as *G. mellonella*, may allow for rapid testing to optimize the approach.

## Abbreviations

CFU: colony-forming-units
K-wires: Kirschner wires
MRSA: methicillin-resistant *Staphylococcus aureus*
PBS: phosphate-buffered saline
PFU: plaque-forming-units
PJIs: Periprosthetic joint infections
SEM: scanning electron microscopy
SM: saline-magnesium buffer
TSA: Tryptic Soy agar
TSB: Tryptic Soy broth

## Declarations

### Ethics approval and consent to participate

Not applicable.

### Consent for publication

All authors have reviewed the manuscript and consent to its publication.

### Availability of data material

The data described in this Research Note can be freely and openly accessed on Figshare. Please see the below table and reference list for details and links to the data:

| Label | Name of data file/data set | File types<br>(file extension) | Data repository and identifier<br>(DOI or accession number) |
| --- | --- | --- | --- |
| Data file 1 | Figure 1 - Galleria Survival Curve | JPG File (.jpg) | Figshare<br>(https://doi.org/10.6084/m9.figshare.30988297) [13] |
| Data file 2 | Galleria Percent Survival Data | MS Word file (.docx) | Figshare<br>(https://doi.org/10.6084/m9.figshare.30988333) [14] |
| Data file 3 | Biofilm K-wire CFUs | MS Word file (.docx) | Figshare<br>(https://doi.org/10.6084/m9.figshare.30988375) [15] |
| Data file 4 | Figure 2 - Biofilm K-wire SEM | PDF file (.pdf) | Figshare<br>(https://doi.org/10.6084/m9.figshare.30988447) [16] |

## Authors’ contributions

T.A., B.W.M.C., and T.J.L. developed and optimized the protocol, and collected and analyzed the data. T.A. wrote and prepared the manuscript for submission. B.W.M.C. and T.J.L. critically reviewed the article. N.T. performed SEM imaging work. S.S.T. and D.A.C. provided funding for this study and were scientific advisors.

## Acknowledgements

We would like to thank Dr. Deborah Court for funding the SEM imaging through the University of Manitoba. We would also like to thank Dr. Matthew Bakker for access to the FEI Talos F200X S/TEM electron microscope through the Manitoba Institute of Materials (University of Manitoba) and support for generating SEM images.

